# B-vitamin requirements and phycosphere interactions for the HAB-forming dinoflagellate, *Pyrodinium bahamense* var. *bahamense*

**DOI:** 10.64898/2025.12.19.695638

**Authors:** Lydia Ruggles Howe, Lauren D’ Amore, Cary Lopez, Sugandha Shankar, Margaret Mars Brisbin

**Affiliations:** College of Marine Science, University of South Florida, St. Petersburg, FL, U.S.A; Florida Fish and Wildlife Conservation Commission, Fish and Wildlife Research Institute, St. Petersburg, FL, U.S.A

**Author notes:** Corresponding Author: 830 1st Street South, St. Petersburg, Florida, U.S.A.

**Keywords:** Cobalamin, dinoflagellate, *Pyrodinium*, Harmful Algal Bloom, HAB, Phycosphere, phytoplankton-bacteria interactions, B-vitamins, B_12_

## Abstract

*Pyrodinium bahamense* is a saxitoxin-producing dinoflagellate found in tropical and subtropical estuarine waters globally, with the var. *bahamense* variety forming recurrent harmful algal blooms (HABs) in Florida estuaries. The intensity and duration of bloom peaks in Florida estuaries vary interannually, but life cycle transitions, estuarine water residence time, water temperature, salinity, and nutrient concentrations appear to contribute to bloom formation. However, the impact of B-vitamin availability on *P. bahamense* bloom formation has not been investigated, despite the knowledge that many HAB dinoflagellates require B-vitamins for growth and bloom development. In this study, we determined that of the three B-vitamins most commonly required by marine phytoplankton—biotin (B_7_), thiamine (B_1_), and cobalamin (B_12_)—*P. bahamense* strictly requires an exogenous B_12_ source to sustain growth. As bacterial interactions can fulfill phytoplankton B_12_ requirements, we tested whether bacterial communities recruited from two Florida estuaries could rescue *P. bahamense* from B_12_ limitation. We demonstrate that bacteria from the Indian River Lagoon (IRL) estuary collected during a *P. bahamense* bloom restored *P. bahamense* growth in media missing B_12_, while bacteria present in Old Tampa Bay (OTB) water collected when *P. bahamense* was not present did not restore *P. bahamense* growth in media missing B_12_. Putative bacterial B_12_ producers exclusively recruited from IRL included Enterobacteraceae genera that likely have anthropogenic sources. Our findings suggest B_12_ is essential for fueling *P. bahamense* blooms, and that specific bacterial communities may contribute to bloom development through B_12_ production.

## Introduction

Phytoplankton are critical to aquatic ecosystem functioning and global biogeochemical cycles [1–3], but phytoplankton can also negatively impact aquatic ecosystems through the formation of harmful algal blooms (HABs). During a HAB, phytoplankton species undergo mass proliferation and may also produce toxins, leading to water column hypoxia, benthic anoxia, wildlife mortalities, and human illnesses [4, 5]. Thus, it is imperative to fully understand the physiology of HAB-forming phytoplankton species to build predictive models for bloom initiation, duration, and intensity. This is increasingly important as changing climate and weather patterns are amplifying the effects of HABs on coastal ecosystems and communities [6].

B-vitamins—such as thiamine (B_1_), biotin (B_7_), and cobalamin (B_12_)—are integral to the growth of many phytoplankton species, as they are essential cofactors for key enzymes involved in cellular metabolism and reproduction [7]. Genomic surveys indicate that more than half of all phytoplankton and up to 90% of HAB dinoflagellates are only equipped with the B_12_-dependent methionine synthase (MetH) [8], making them fully reliant on external B_12_ sources for essential amino acid synthesis. Through a comprehensive set of bioassays, Tang et al. (2012) demonstrated that 26 out of 27 HAB species (41 strains) required B_12_ and 20 required B_1_ [9], further suggesting B-vitamin availability is especially critical to the growth of HAB-forming phytoplankton. However, the extent of the role B-vitamin availability plays in promoting or limiting HABs is not fully elucidated [10].

Despite reliance on B vitamins, many HAB species bloom in coastal waters where measurable dissolved B-vitamin concentrations are below phytoplankton growth requirements, suggesting that B vitamins may be limiting nutrients in coastal estuaries [10]. Moreover, studies in Long Island estuaries demonstrated a drawdown of vitamins B_1_ and B_12_ during dinoflagellate blooms, and B_1_ and B_12_ amendments in concurrent incubation experiments increased dinoflagellate biomass [11]. Bacterial production of B vitamins stands to be the major source of these critical resources to marine ecosystems. Roughly 25% of sequenced bacterial genomes contain complete B_12_-synthesis pathways [12], but most B_12_ producers do not freely share the resource [13]. Thus, specific relationships between phytoplankton and bacteria inhabiting the phycosphere may be particularly important for fulfilling phytoplankton B_12_ requirements [14, 15]. Indeed, positive interactions between several HAB dinoflagellates and specific B-vitamin producing bacteria have been identified [16–18].

The saxitoxin-producing HAB dinoflagellate, *Pyrodinium bahamense* var. *bahamense,* forms dense assemblages (> 100,000 cells L^-1^) in Florida estuaries, including in the northern segment of Tampa Bay, Old Tampa Bay (OTB), on Florida’s west coast and the Indian River Lagoon (IRL) estuary on the east coast [19] (Fig. 1). B vitamins have long been hypothesized to be contributing factors in supporting the high concentrations of *P. bahamense* observed in the bioluminescent bays of Puerto Rico, where the dinoflagellate blooms carry ecological, cultural, and economic significance [20]. However, the role of B-vitamin availability and bacterial interactions in supporting *P. bahamense* blooms in Florida estuaries has yet to be directly investigated, making it plausible that characterizing these interactions may improve our understanding of bloom intensity and duration in the region. This study directly investigates the B-vitamin requirements of a *P. bahamense* strain isolated from OTB and determines whether bacterial communities recruited from OTB and IRL seawater can support *P. bahamens*e growth in the absence of vitamin B_12_.

**Fig. 1.**
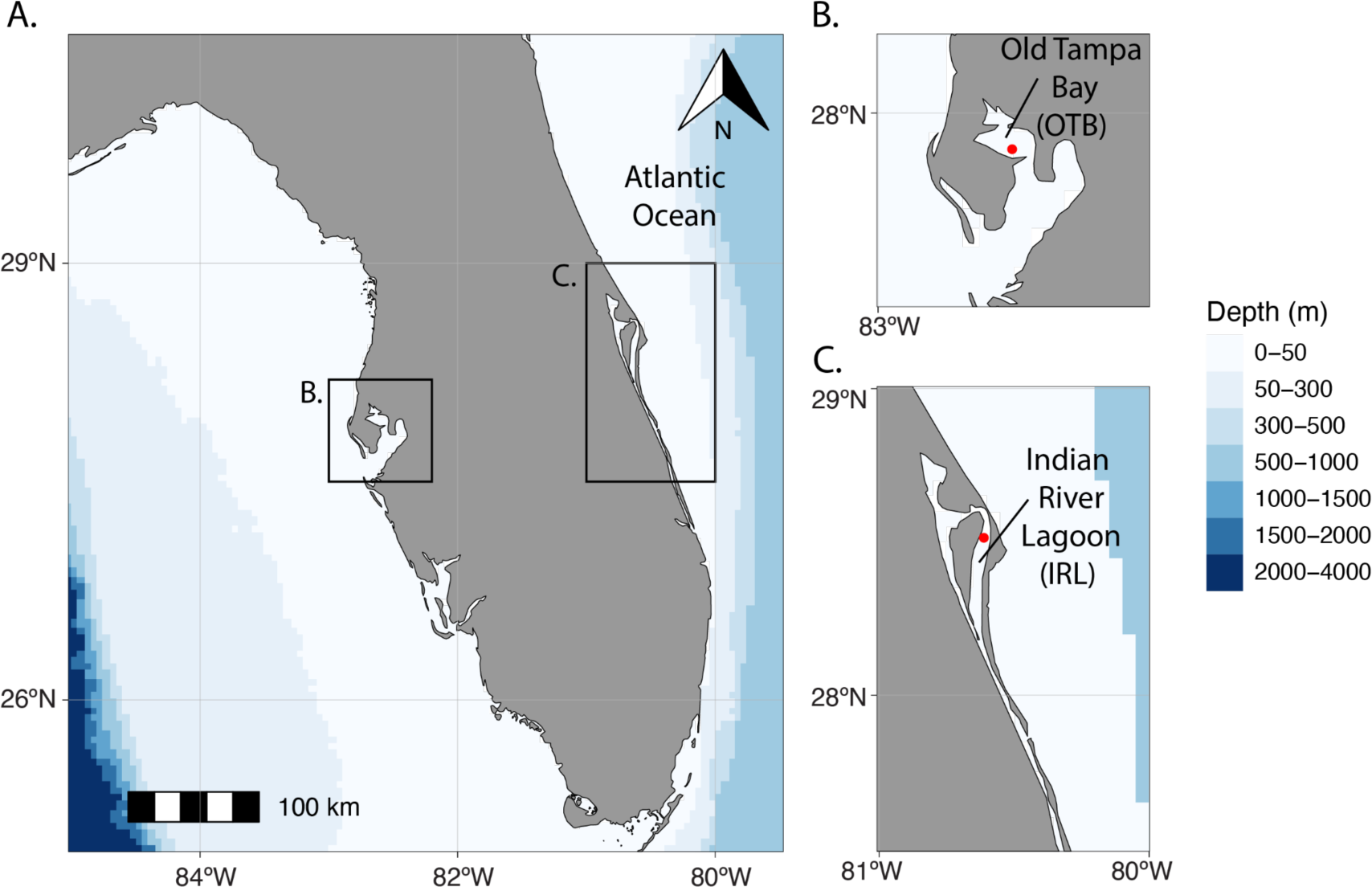
Map depicting regions of Florida (U.S.A.) with recurrent *Pyrodinium bahamense* blooms and where samples were collected for this study. **A** Map of Florida with inset regions indicated (Tampa Bay and OTB; **B** and Indian River Lagoon; **C**). Red points in panels **B** and **C** indicate the locations where seawater was collected for bacterial community transfers to FWC1006.

## Materials and Methods

### P. bahamense *strains used in experiments*

*P. bahamense* strains FWC1006—originating from OTB in 2023—and FWC1003—originating from the IRL in 2017—were acquired from the Florida Fish and Wildlife Conservation Commission (FWC) Fish and Wildlife Research Institute (FWRI) for use in this study (Fig. 1). Both strains were established by germinating resting cysts from sediment samples. Germinated cells were then isolated and grown into unialgal cultures using aseptic techniques, but were not deliberately rendered axenic. Upon transfer to our lab, both strains were maintained in L1/4 media prepared with sterile 25 PSU artificial seawater (ASW; Instant Ocean) [21] at 24°C on a 12:12 h light: dark cycle with 50% illumination (∼115 µM photons m^-2^ s^-1^) in a Percival I-36 LLVL incubator.

### *Characterization of bacterial communities in* P. bahamense *cultures*

Bacteria associated with phytoplankton can provide B vitamins to support phytoplankton [7, 16, 22], making it imperative to characterize bacteria co-cultured with *P. bahamense* before experimentation. The presence of bacteria in *P. bahamense* strains FWC1006 and FWC1003 was initially evaluated by transferring aliquots into two test broths for bacterial growth: Difco Marine Broth and an artificial seawater tryptone broth (ASWT) matching the salinity of algal culture media [23]. ASWT broth was prepared from 1 L of 25 PSU Instant Ocean ASW amended with 5 g tryptone, 3 g yeast extract, and 3 mL 50% glycerol (in water, RICCA Chemical Company), and sterilized by autoclaving and filtering with a 0.2-µm pore-size filter [23]. Aliquots of each isolate (100 µl) were transferred to triplicate wells containing 1 mL of test broth in a sterile 24-well plate. Triplicate negative controls for each test broth were included. Plates were incubated at room temperature, and bacterial growth was visually monitored after 72 h and checked under 400 x magnification with an inverted microscope.

Bacterial communities present in cultures of the two *P. bahamense* strains were further characterized through metabarcode sequencing. Biomass was collected from each strain by sequentially filtering cultures through 10.0-µm and 0.2-µm pore-size filters under gentle vacuum pressure. *P. bahamense* cells and associated bacteria were retained on 10.0-µm pore-size filters, and free-living bacteria were retained on 0.2-µm pore-size filters. RNA was extracted from filters using the RNeasy PowerWater Kit (Qiagen) according to the manufacturer’s protocol with the suggested 10-minute 65°C incubation for improved cell lysis. RNA was reverse transcribed following the manufacturer’s protocols for the High-Capacity RNA-to-cDNA Kit (Applied Biosystems). The V3–V4 hypervariable regions of the reverse-transcribed 16S rRNA were amplified following the “16S Metagenomic Sequencing Library Preparation” Illumina protocol, including the recommendations for primers [24] and PCR conditions. Metabarcode libraries were sequenced on an Illumina NextSeq2000 2x300-bp P1 flow cell. Amplicons were processed with the QIIME2 pipeline [25], utilizing the DADA2 denoising algorithm [26], and representative sequences were taxonomically classified using a Naive Bayes classifier [27] trained on the SILVA Ref 99 database v138.1 [28]. Amplicon Sequence Variants (ASVs) were further processed using the phyloseq [29] package in R Studio [30]. Based on results from bacterial growth tests and metabarcode analyses, the FWC1006 *P. bahamense* isolate from OTB was selected for subsequent experiments.

### *Evaluating the B-vitamin requirements of* P. bahamense

Given the difficulty of growing dinoflagellates axenically, dinoflagellate cultures with reduced or knocked-down microbiomes have been used in experiments to minimize the impact of bacterial processes on experimental results [17]. Here, we propose that the low-diversity bacterial community associated with the FWC1006 *P. bahamense* strain from OTB represents a reduced bacterial community and an opportunity for testing the B-vitamin requirements of *P. bahamense* without subjecting cultures to extensive antibiotic treatment.

To determine the B-vitamin requirements for *P. bahamense*, the OTB strain was grown in L1/4 media missing one of the three major B-vitamins—B_12_ (cobalamin), B_1_ (thiamine), or B_7_ (biotin)—and in replete L1/4 (i.e., containing all three B vitamins) as a positive control. All media formulations were made using sterile 25 PSU ASW amended with NaNO_3_, NaH_2_PO_4_, and trace metal solution from the NCMA L1 Medium kit at 1/4 the concentration of L1 medium [21]. Individual B vitamins (Sigma-Aldrich) were added to media at 1/4 the concentration in L1 medium [21]. Experimental cultures were set up with 1 mL of the OTB isolate and 45 mL of each media treatment in triplicate 50 mL cell culture flasks and incubated as described above [31]. Growth was monitored by measuring *in vivo* Chlorophyll-*a* fluorescence (RFU) with a Trilogy fluorometer (Turner Designs) every 48 h. The experiment was maintained for four trials, with 1 mL serial transfers made in the late exponential growth phase or early stationary phase. The – B_12_ treatment was not serially transferred because these cultures failed to grow. Instead, 1 mL of the culture in replete media was also transferred to –B_12_ media at the beginning of each new trial. Growth rates were calculated from Chlorophyll-*a* fluorescence using the fit_easylinear function from the growthrates R package (h = 5, quota = 1) [32].

To better characterize the minimum B_12_ requirement of *P. bahamense*, the OTB isolate was grown in L1/4 media with varying final concentrations of vitamin B_12_ from 369 pM—the concentration in L1 medium [21]—through 0.0369 pM, prepared through 10% serial dilutions. Culture conditions matched those described above, and *in vivo* Chlorophyll-*a* fluorescence was measured every 48 h for a single 40-day trial.

### *Evaluation of bacterial community potential to supplement B_12_ and support* P. bahamense *blooms*

Next, we tested the ability of bacterial communities from OTB and IRL to produce vitamin B_12_ and promote *P. bahamense* growth. OTB surface seawater was collected on May 2, 2024 (27.9055°N, –82.6046°W) and IRL surface seawater was collected on July 10, 2024 (28.5136°N, –80.6125°W) by FWRI (Fig. 1). At the time of collection, no *P. bahamense* cells were detected at the OTB site, whereas *P. bahamense* cells were detected at concentrations exceeding the FWC-designated bloom level at the IRL site (113,333 cells L^-1^, bloom-level: >100,000 cells L^-1^). Seawater from both sites and a *P. bahamense* culture (IRL FWC1003) were sequentially filtered through 10.0-µm, 0.8-µm, and 0.45-µm pore-size filters to remove eukaryotic phytoplankton, but particle-associated bacteria and larger or filamentous bacteria were likely also removed or reduced during filtering. A 250 mL aliquot of the final filtrate for each recruitment treatment was filtered through a 0.2-µm pore-size filter, and the remaining filtrate for each recruitment treatment was diluted 50% with L1/4 media and used in recruitment trials. The 10.0-µm and 0.2-µm filters were flash frozen and stored at –80°C for DNA extraction.

A 1-mL aliquot of OTB *P. bahamense* culture was transferred into 45 mL of each of the three recruitment treatment filtrates (OTB seawater, IRL seawater, IRL culture) and incubated for two weeks to allow equilibration of the introduced bacterial communities. Subsequently, 1 mL aliquots of cultures with recruited bacterial communities were transferred in triplicate to 45 mL of replete L1/4 and media without B_12_ (L1/4 –B_12_), and grown and monitored as described above for two trials. For the second trial, cultures were serially transferred if growth occurred, which was the case for all cultures grown in replete media but only for the IRL seawater-treated culture grown in L1/4 –B_12_. The OTB seawater- and IRL culture-treated cultures in L1/4 –B_12_ were restarted from the matching treatment in replete media in trial 2, since they failed to grow in trial 1. All culture replicates from growth trial 2 were sequentially filtered through 10.0-µm and 0.2-µm pore-size filters under gentle vacuum pressure when cultures began entering stationary growth phase. Filters were flash frozen and stored at –80°C for DNA extraction.

### Nanopore full-length 16S metabarcoding and bioinformatics

The bacterial communities introduced from seawater sources to OTB *P. bahamense* and the established “recruited” communities from each source (IRL *P. bahamense* culture, IRL seawater, and OTB seawater) were assessed through full-length 16S rRNA gene sequencing on the Oxford Nanopore MinION platform. DNA was extracted from 10.0-µm and 0.2-µm pore-sized filters with the Qiagen DNeasy PowerSoil Pro kit according to the manufacturer’s protocol, including the additional 65°C heating step. The full-length 16S rRNA gene was amplified using forward primer S-D-008-c-S-20 (5′-AGRGTTYGATYMTGGCTCAG-3′) [24] and reverse primer 1492R (5′-RGYTACCTTGTTACGACTT-3′) [33] with KAPA2G Robust HotStart (Roche) in the following PCR program: 95°C/ 3min, 35x (95°C/ 20sec, 57°C/ 20sec, 72°C/ 30sec), 72°C/ 1min, and 4°C/ ∞ [33]. Sequencing libraries were prepared using the Oxford Nanopore Native Barcoding Kit 24 v14 following the manufacturer’s protocol and sequenced on two MinION v10.4 flow cells and one Flongle flow cell. Sequence data were processed (min. length = 1300 bp, max. length = 2000 bp), quantified, and classified using Kraken v2.1.2 [34] within the wf-16s nextflow pipeline in the EPI2ME platform [35] with the SILVA v138.1 taxonomic database [28]. Results were exported to R Studio for further processing.

## Results

### *Assessment of bacterial communities co-cultured with* P. bahamense *isolates*

There was no visible bacterial growth (broth turbidity or observable cells by microscopy) in Difco Marine Broth or ASWT broth 72 h after inoculation with FWC1006 *P. bahamense* culture. Similarly, there was no visible bacterial growth in Difco Marine Broth or in salinity-matched ASWT broth in the negative control wells after the 72-hour observation period. In contrast, there was visible bacterial growth (high turbidity) in both Difco Marine Broth and salinity-matched ASWT broth in wells inoculated with FWC1003 *P. bahamense* culture. Further investigation into the bacterial community composition in cultures of both *P. bahamense* strains through metabarcoding also revealed stark differences. The bacterial community in the OTB *P. bahamense* culture had very low alpha diversity (Shannon Index) compared to the communities in IRL *P. bahamense* cultures (Fig. 2A). The IRL *P. bahamense* culture hosted bacteria from 11 taxonomic orders, including typical phytoplankton-associated groups such as Alteromonadales, Caulobacterales, and Rhodobacterales (Fig. 2B). Conversely, the OTB *P. bahamense* culture hosted bacteria from just three orders: Microtrichales, Oceanospirillales, and Rhizobiales (Fig. 2B). At the amplicon sequence variant (ASV) level, only nine ASVs were detected among OTB *P. bahamense* cultures and just six ASVs made up > 99% of sequencing reads (Table S1). Further investigation of these ASVs through BLAST searches revealed that they were previously found in soils, marine sediments, and aquatic plant rhizospheres (Table S1). Thus, the OTB *P. bahamense* culture has a reduced bacterial community compared to the IRL *P. bahamense* culture. Moreover, the bacteria associated with the OTB *P. bahamense* culture likely do not represent water column bacteria that *P. bahamense* would interact with during blooms of vegetative cells.

**Fig. 2.**
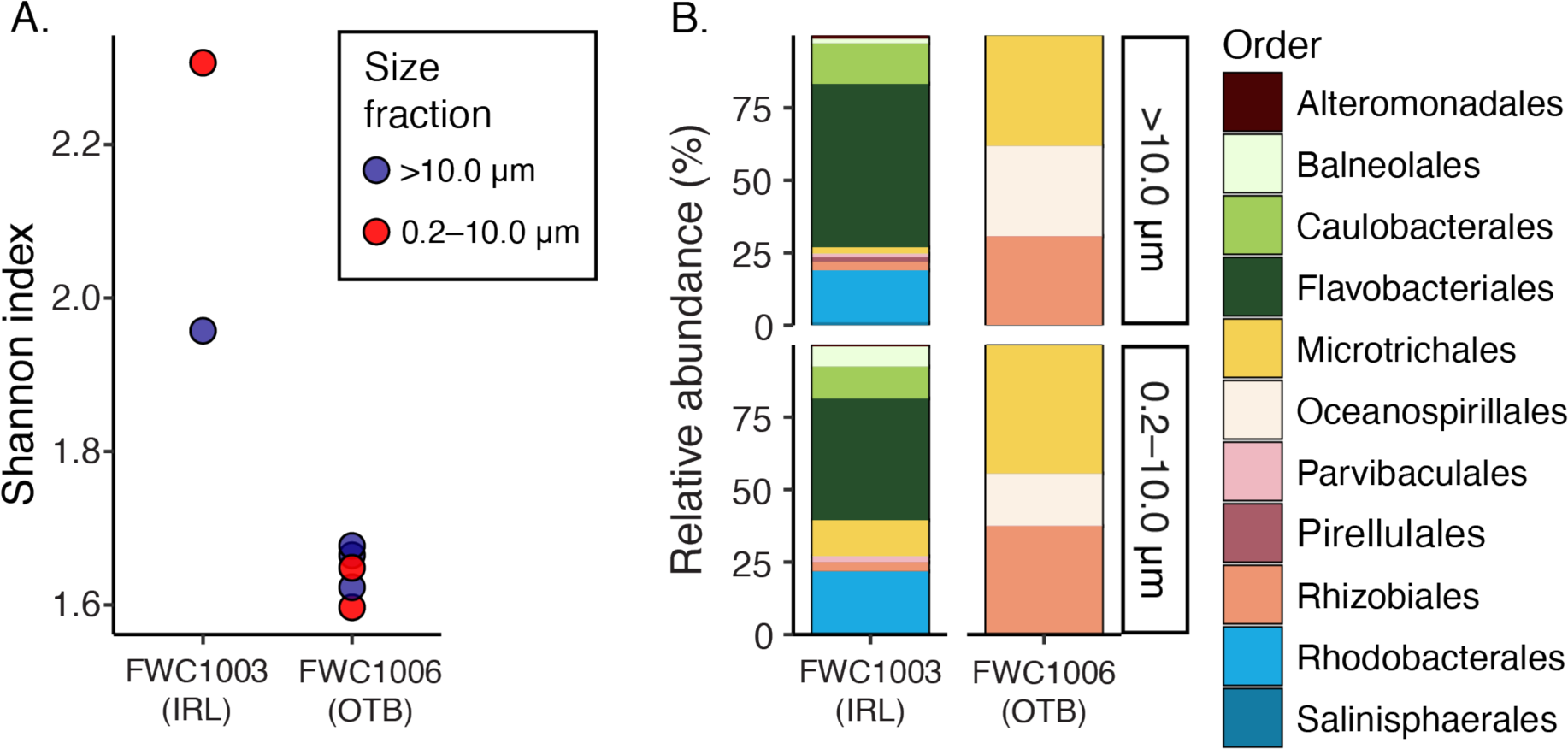
Alpha diversity and composition of bacterial communities in *P. bahamense* cultures. **A** Shannon diversity index for particle-associated (>10.0-µm size fraction; blue) and free-living (0.2–10.0-µm size fraction; red) bacterial communities in cultures of FWC1003 and FWC1006 *P. bahamense* isolated from Indian River Lagoon and Old Tampa Bay, respectively. **B** Relative abundance of bacterial orders in size-fractioned communities in *P. bahamense* cultures.

### *B-vitamin requirements of OTB* P. bahamense

Given the low-diversity bacterial community detected in the OTB *P. bahamense* culture, this strain was used in experiments testing the B-vitamin requirements of *P. bahamense*. We hypothesized that the low-diversity sediment-derived bacteria in the OTB culture would be less likely to symbiotically provide B-vitamins than the higher-diversity community observed in the IRL culture, which was more similar to communities observed in other phytoplankton cultures (e.g., [36–38]).

Growth curves for OTB *P. bahamense* appeared similar across four serial transfers in replete L1/4 media, L1/4 –thiamine, and L1/4 –biotin (Fig. 3A). In contrast, OTB *P. bahamense* immediately ceased growing when transferred into L1/4 media missing vitamin B_12_ (L1/4 –B_12,_ Fig. 3A). As a result, serial transfers could not be made for this treatment, and culture from the replete treatment was transferred to L1/4 –B_12_ in each transfer round. Results from a one-way ANOVA indicated that the effect of media formulation on maximum chlorophyll-*a* fluorescence, a proxy for biomass, was significant (*p* = .0000, *F* = 45.35), and a Tukey HSD multiple comparisons of means test showed that the maximum fluorescence was significantly lower in L1/4 –B_12_ media than in replete (*p-adj* = .0000), –thiamine (*p-adj* = .0000), and –biotin media (*p-adj* = .0000). Of note, the maximum fluorescence reached in media missing thiamine was significantly greater than in replete media (*p-adj* = .0000)or in –biotin media (*p-adj* = 0.0006). Since *P. bahamense* ceased growing when transferred into media missing B_12_, reliable growth rates could not be determined for this treatment. Aside from the –B_12_ treatment, media formulation did not have a large effect on growth rate (Fig. 3C, S1); a one–way ANOVA excluding the –B_12_ treatment indicated that media formulation (i.e., –thiamine, –biotin, replete) did not significantly affect growth rate (*µ*; *p* = 0.172, *F* = 1.861).

**Fig. 3.**
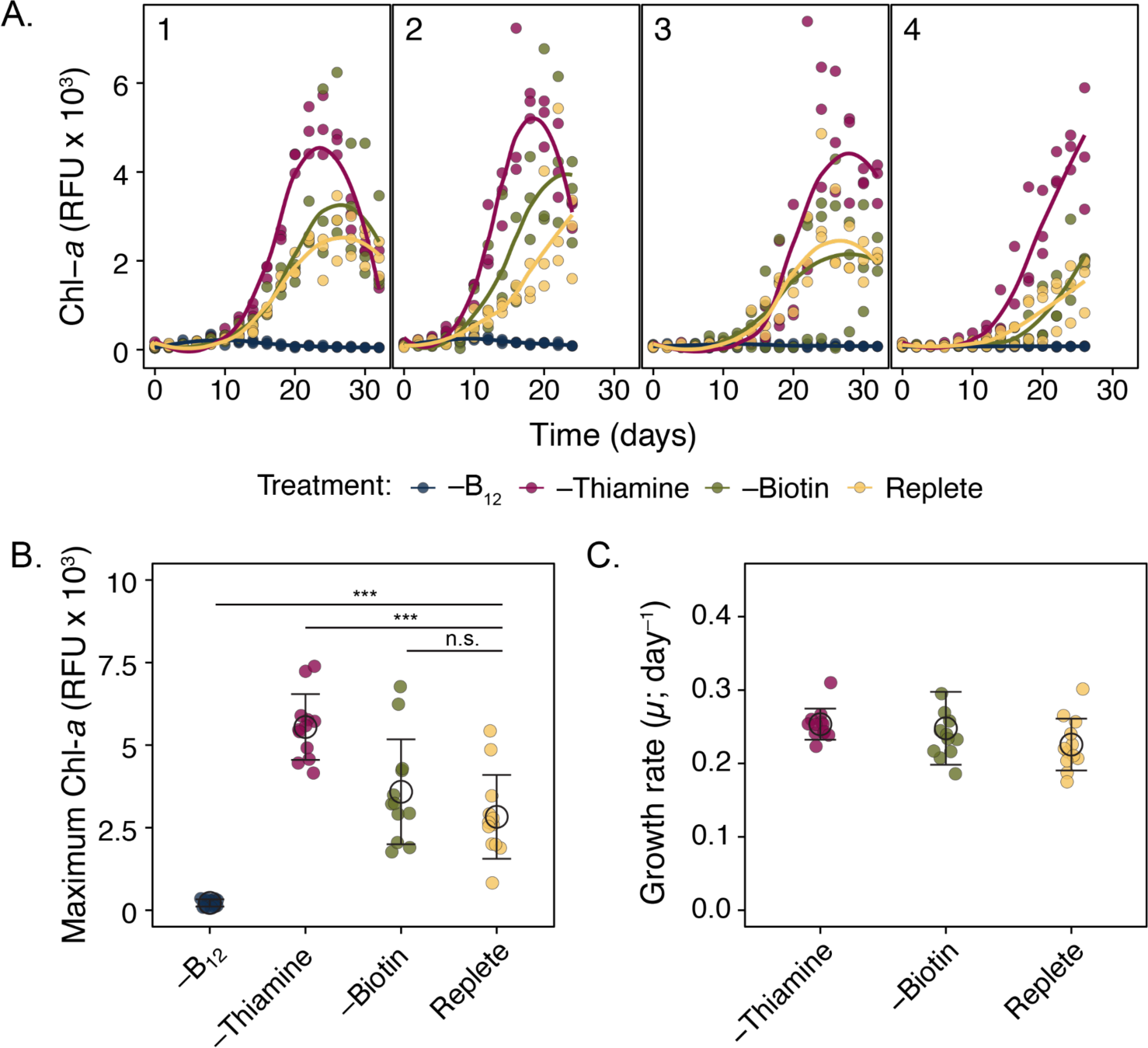
Growth responses of *P. bahamense* when grown in replete L1/4 media and in media missing B_12_, thiamine, or biotin. All data are plotted with color representing media treatment. **A** *In vivo* Chlorophyll *a* fluorescence (Chl-*a*) was used to track *P. bahamense* growth across four trials. Solid lines indicate the smoothed conditional mean, plotted with the geom_smooth function in the R package ggplot2 (method = ‘loess’). **B** Maximum Chl-*a* values in each replicate by media formulation. Results from post-hoc comparisons to replete are shown with the following significance codes: *p-adj* > 0.05; ‘n.s.’ and *p-adj* < 0.001; ‘***’. **C** Growth rate of each replicate in –thiamine, –biotin, and replete media. Growth rates could not be calculated for replicates in –B_12_ media due to the lack of growth (Fig. S1). Means are plotted as black open circles, and error bars represent one standard deviation of the mean.

### P. bahamense *growth response to varying B_12_ availability*

Given that withholding B_12_ rapidly diminished *P. bahamense* growth (Fig. 3), we further evaluated the *P. bahamense* growth response to varying B_12_ concentrations using 10% serial dilutions of B_12_ in L1/4 media. *P. bahamense* growth showed a typical dose-response to increasing B_12_ availability (Fig. 4A). Maximum biomass (Chl-*a* fluorescence) increased with increasing B_12_ availability (Fig. 4B), with significant increases in maximum fluorescence between 0.0369 and 3.69 pM B_12_ and between 3.69 and 36.9 pM B_12_ (*p-adj* = 0.025 and 0.0000, respectively; Tukey HSD). However, when enough B_12_ was available to support growth, the growth rates were fairly consistent (Fig. 4C, S2). Growth rate was depressed in media with 0.0369 pM B_12_ (Fig 4C), but one-way ANOVA results did not indicate that B_12_ concentration significantly affected growth rate (*p* = 0.196, *F* = 1.85, Fig 4C). Notably, Chl-*a* fluorescence in 0.0369 pM B_12_ media remained low and variable throughout the trial, leading to poor linear model fits for estimating growth rate (*p* > 0.05, *R*^2^ < 0.85, Fig. S2) that appear to overestimate growth rates and skew the statistical test results.

**Fig. 4.**
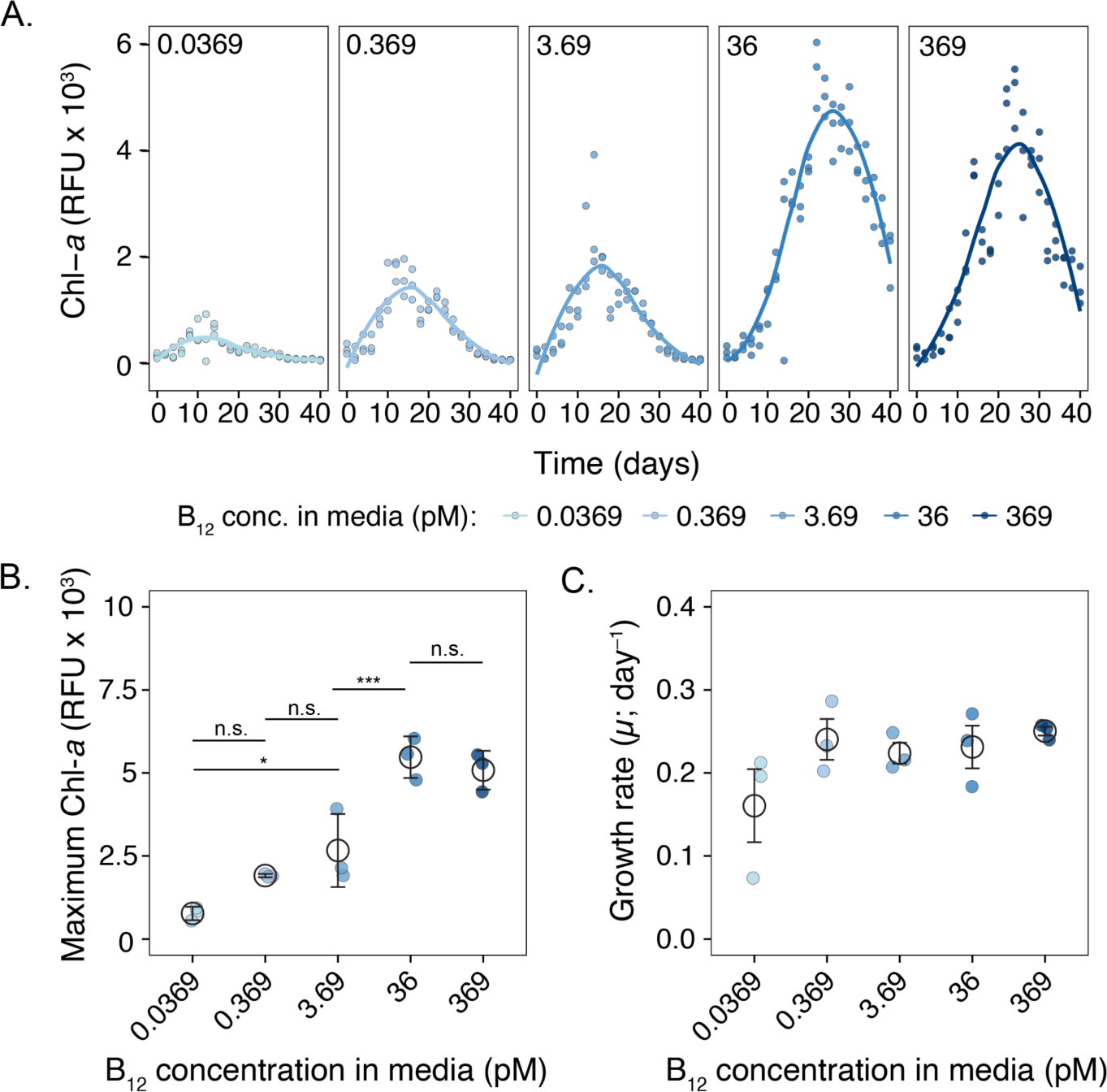
Growth responses of *P. bahamense* when grown in L1/4 media with a 10% dilution series of B_12_ concentrations. All data are plotted with color representing the B_12_ concentration. *In vivo* Chlorophyll-*a* fluorescence (filled circles) was used to track *P. bahamense* growth in L1/4 media with five concentrations of B_12_: 0.0369 pM, 0.369 pM, 36 pM, and 369 pM. Solid lines indicate the smoothed conditional mean, plotted with the geom_smooth function in the R package ggplot2 (method = ‘loess’). **B** Maximum Chl-*a* fluorescence values and **C** growth rates (filled circles), shown for each replicate grouped by media B_12_ concentration. Means are plotted as black open circles, and error bars represent one standard deviation of the mean. Results from key post-hoc comparisons are shown with the following significance codes: *p-adj* > 0.05; ‘n.s.’, *p-adj* < 0.05; ‘*’, and *p-adj* < 0.001; ‘***’.

### *Effects of recruited bacterial communities on* P. bahamense *growth*

Increased growth was detected in OTB *P. bahamense* (FWC1006) with bacteria recruited from IRL seawater compared to *P. bahamense* with bacteria recruited from IRL culture or OTB seawater when grown in L1/4 –B_12_ media (Fig. 5A). One-way ANOVA indicated that the recruited bacterial community significantly affected maximum Chl-*a* when B_12_ was withheld (Fig. 5B; *p* = 0.0001, *F* = 50.27). The maximum Chl-*a* in cultures with bacteria from IRL seawater was significantly higher than in cultures with bacteria from OTB seawater, bacteria from the IRL culture, or with the initial bacteria in the OTB culture (*p-adj* = 0.0000 for each comparison, Tukey HSD). Growth rates for *P. bahamense* with bacteria from IRL seawater were not significantly different when grown in replete media versus media missing B_12_ (two-sample t-test; *p* = 0.5678, *t*(10) = –0.5907; Fig. 5C, S3). These results indicate that at least some of the bacteria in the community recruited from IRL seawater are B_12_ producers and can support P. *bahamense* growth.

**Fig. 5.**
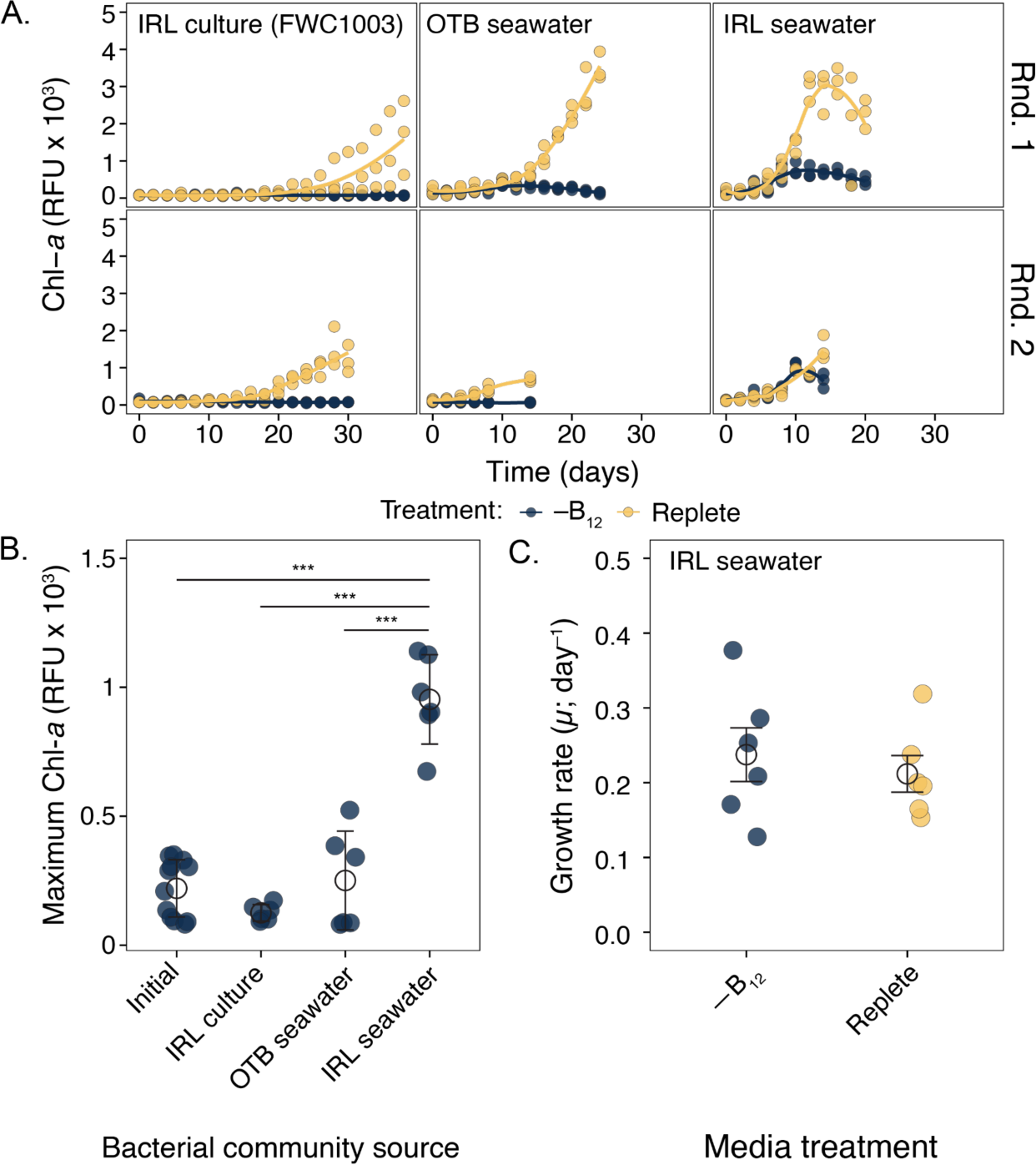
Growth responses of OTB *P. bahamense* (FWC1006) with bacterial communities recruited from IRL culture (FWC1003), OTB seawater, and IRL seawater when grown in L1/4 media missing vitamin B_12_ and replete L1/4 media. All data are plotted with color representing media treatment. **A** *In vivo* Chlorophyll-*a* fluorescence (filled circles) was used to track the growth of OTB *P. bahamense* with bacteria from three sources (IRL culture, OTB seawater, and IRL seawater) in replete L1/4 media and L1/4 media missing B_12_ (L1/4 –B_12_). Solid lines indicate the smoothed conditional mean, plotted with the geom_smooth function in the R package ggplot2 (method = ‘loess’). **B** Maximum Chl-*a* fluorescence values (filled circles) in L1/4 –B_12_ are shown for each replicate, grouped by bacterial community source. Note, the maximum Chl-*a* reached in L1/4 –B_12_ by OTB *P. bahamense* with its initial low-diversity bacterial community is included for reference (from Fig. 3). Results from key post-hoc comparisons are shown with the following significance code: *p-adj* < 0.001; ‘***’. **C** Growth rates (filled circles) for each replicate with bacteria from IRL seawater, grouped by media treatment. Means are plotted as black open circles, and error bars represent one standard deviation of the mean.

### Sequencing results

Overall, this project produced 10,431,416 Nanopore full-length 16S reads that passed quality filters. After applying a length filter (min = 1300 bp, max = 2000 bp), 8,526,020 reads remained. Culture samples sequenced on MinION flow cells yielded 128,931–500,307 (median = 322,769) reads per sample and retained 103,950–407,502 (median = 272,432) after applying the length filter. Seawater samples (bacterial source cultures) sequenced on a Flongle flowcell yielded 34,389–141,236 (median = 93,931) reads per sample and retained 12,221–53,990 (median = 37,987) reads per sample after length filtering. Alpha diversity (richness and Shannon index, based on genus) was higher in the IRL seawater (richness = 1,695–1,805, Shannon = 4.82–4.89) than the OTB seawater (richness = 1,056–1,208, Shannon = 3.24–4.69; Fig. S5, S6). Alpha diversity was similar across recruited communities (Fig. S5, S6).

### Comparison of bacterial communities recruited from different sources

Distinct bacterial communities were recruited to the FWC1006 *P. bahamense* cultures from the three sources (FWC1003 *P. bahamense* cultures (IRL), IRL seawater, and OTB seawater). Recruited bacterial communities formed three clusters in a principal coordinate analysis based on Bray-Curtis distances (Fig. 6). PERMANOVA results further indicated that bacterial source significantly affected recruited community composition (*R*^2^ = 0.67542, *F* = 16.127, *p* = 0.001; 999 permutations), whereas size fraction and media formulation did not significantly affect community composition (*p* > 0.05). Visualizing the relative abundance of bacterial orders in each sample further demonstrates the distinct communities recruited from each source (Fig. 7). Moreover, the FWC1006 *P. bahamense* culture clearly recruited bacteria from the various sources; only three bacterial orders were detected in the initial FWC1006 culture microbiomes, despite deep Illumina metabarcode sequencing (394,399–621,150 reads per sample after all filterings steps), whereas 10–12 bacterial orders were detected after recruiting new bacterial communities.

**Fig. 6.**
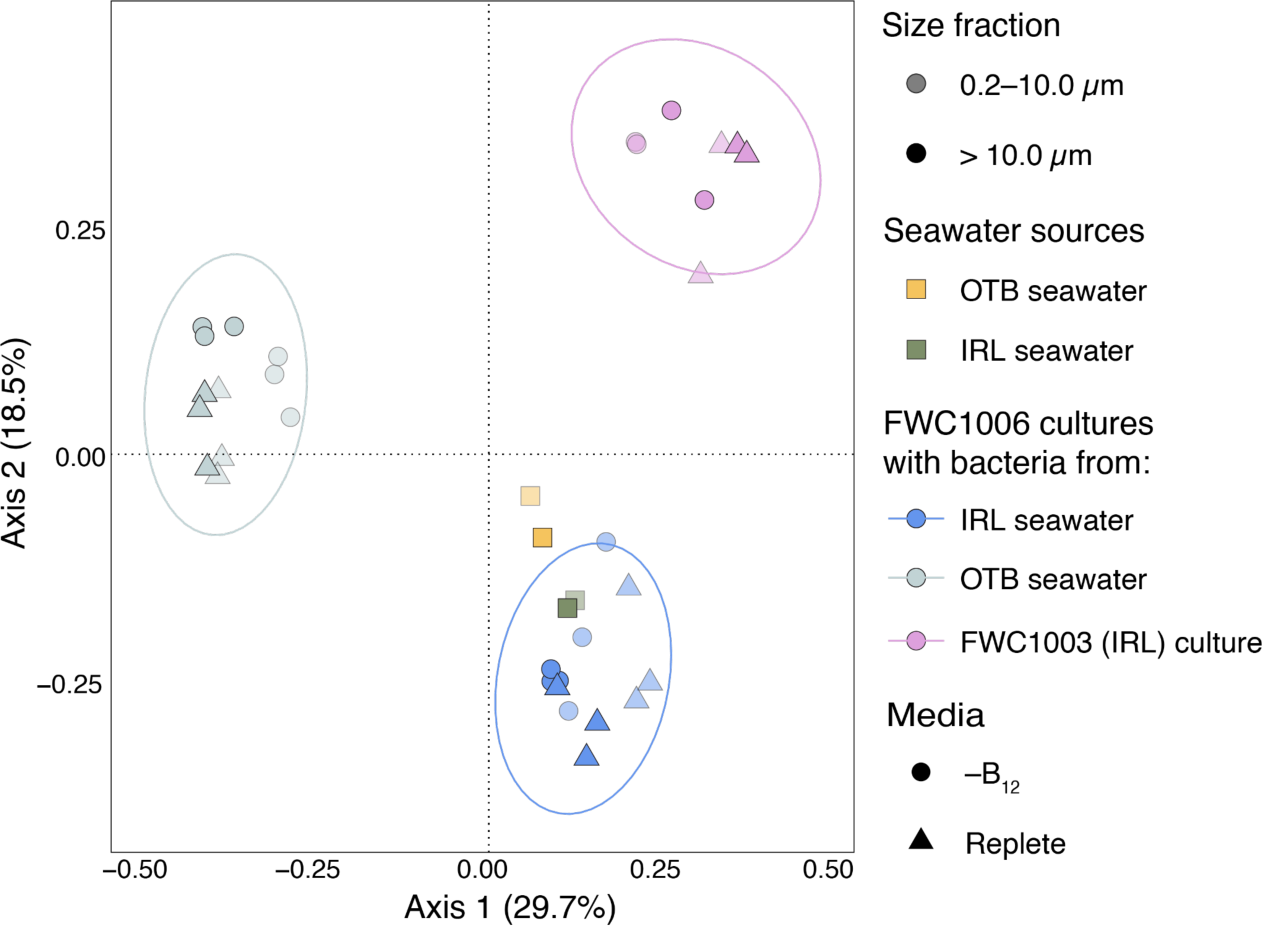
Principal coordinate analysis (PCoA) of Bray-Curtis distances between the bacterial communities recruited by *P. bahamense* FWC1006 and the source surface seawater bacterial communities. FWC1006 cultures with each bacterial community in replete L1/4 media and L1/4–B_12_ media were sacrificed for 16S rRNA gene sequencing at the end of round two in Fig. 5C. Three 95% confidence ellipses drawn with the ggplot2 stat_ellipse function highlight the groupings within the data. The replicates with bacteria from OTB seawater separate from all other samples on the primary axis, while the samples with bacteria from the FWC1003 (IRL) culture and IRL seawater separate on the secondary axis. The bacterial communities in the source seawater grouped most closely with the bacterial communities in culture that were recruited from IRL seawater (bottom right quadrant), with the IRL seawater source communities within the 95% confidence ellipse for the communities recruited from IRL seawater.

**Fig. 7.**
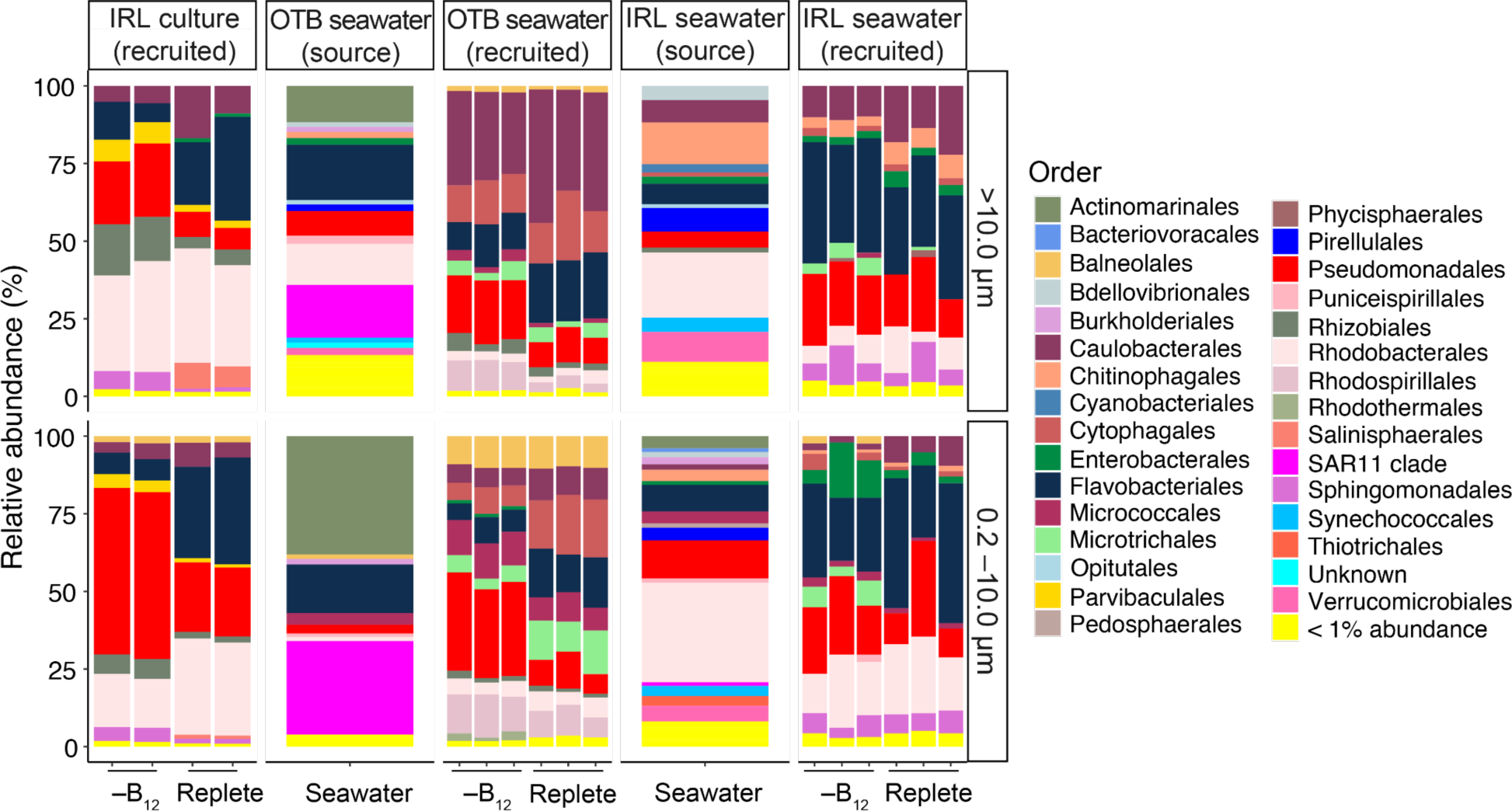
Relative abundance of bacterial orders in source seawater samples and in cultures with bacterial communities recruited from IRL P. bahamense culture, OTB seawater, and IRL seawater. Orders that made up less than 1% in a given sample were combined to reduce the number of orders included in the plot.

Since only cultures with bacteria recruited from IRL seawater were able to grow in media without vitamin B_12_, we identified bacteria that were exclusively present in these cultures. Alteromonadaceae, Saprospiraceae, and Enterobacteriaceae were consistently present in communities recruited from IRL seawater but not from OTB seawater or IRL culture (Fig. 8). *Alteromonas spp*., which typically require B_12_ but can not synthesize it [39], were abundant in the smaller size fraction of communities recruited from IRL seawater when B_12_ was not included in the media. *Klebsiella* and *Enterobacter*—both B_12_ producers [40, 41]—were present in all communities recruited from the IRL seawater but not from OTB seawater or IRL culture. Similarly, *Lewinella* was present in all communities recruited from IRL seawater but not from OTB seawater or IRL culture. In contrast, *Ruegeria* and *Sulfitobacter* (Rhodobacteraceae), genera often detected in phytoplankton cultures, were present in all cultures (Fig. 8).

**Fig. 8.**
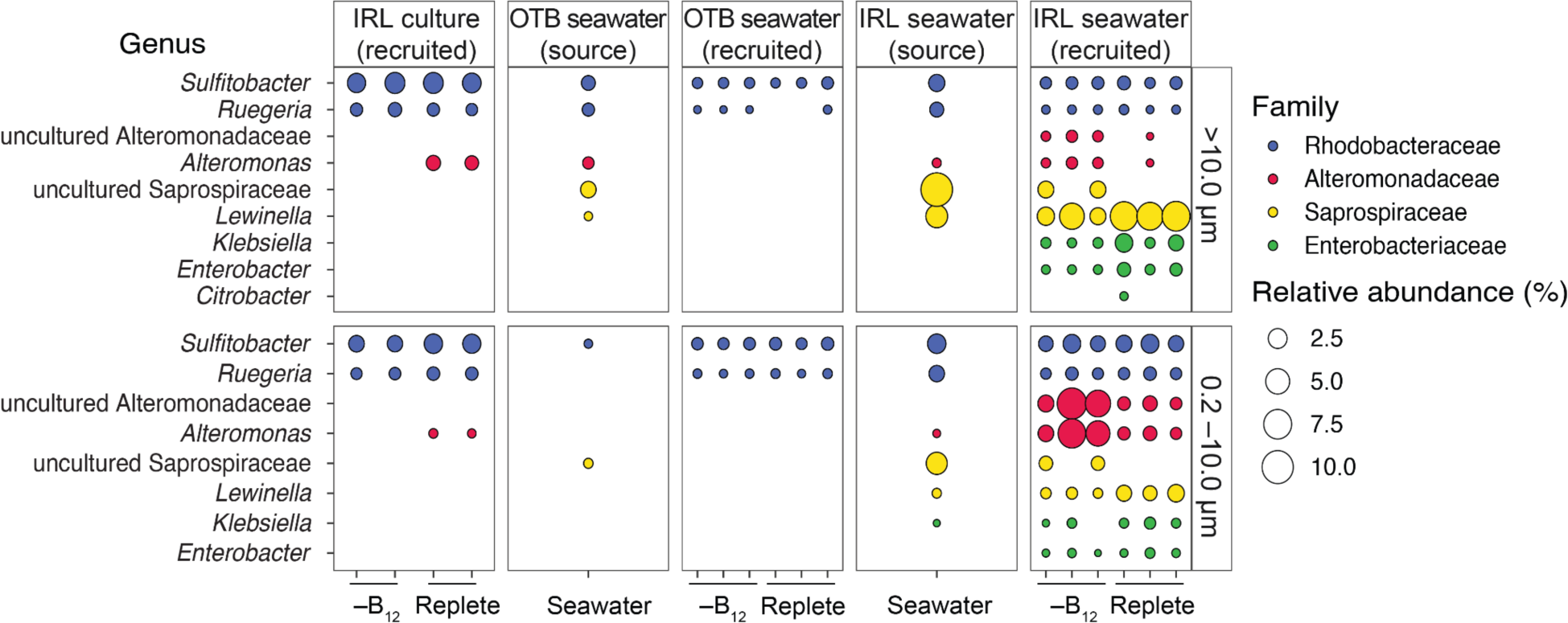
Relative abundance of genera in groups detected primarily in communities recruited from IRL seawater (Alteromonadaceae, Saprospiracea, and Enterobacteracea). Rhodobacteracea is included as a comparison. Rhodobacteracea are commonly found in association with phytoplankton and other algae. *Ruegeria* and *Sulfitobacter* were detected in all sample types.

## Discussion

HAB-forming dinoflagellates generally require B_12_ for growth and often also require B_1_ or B_7_ [9]. Despite this, HABs frequently form in regions with undetectable B_12_ concentrations, suggesting that B_12_ may be being drawn down below detection limits or that cryptic B_12_ cycling from bacteria is key to supporting HABs [10]. Here, we exclusively tested the cobalamin (B_12_), thiamine (B_1_), and biotin (B_7_) requirements of *P. bahamense* var. b*ahamense*—a dinoflagellate that causes toxic HABs in Florida estuaries [19], but whose bioluminescence makes it culturally and economically important in the Caribbean [42]—by utilizing a strain isolated from Florida (OTB) with a reduced culture microbiome (Fig. 1). Our results clearly demonstrate that *P. bahamense* has an absolute B_12_ requirement for growth and bloom formation. Only the bacterial community recruited from IRL seawater collected during a *P. bahamense* bloom restored *P. bahamense* growth in media without B_12_. These results suggest that bacteria present during *P. bahamense* blooms may contribute to fueling blooms through B_12_ production. While bacterial communities that restored P. bahamense growth (from IRL seawater) when B_12_ was withheld overlapped in composition with those that did not (IRL culture and OTB seawater), several specific bacterial genera were found exclusively in the growth-restoring communities. Among these, *Klebsiella* and *Enterobacter* are known B_12_ producers [41, 43] and may be introduced to the environment through wastewater or runoff [44, 45]. Thus, anthropogenic inputs may be contributing to *P. bahamense* blooms in Florida estuaries by introducing or otherwise supporting B_12_-synthesizing bacteria.

### B-vitamin requirements of toxigenic dinoflagellates

Results from this study demonstrate that *P. bahamense* from OTB requires an external source of vitamin B_12_, but not necessarily B_1_ or B_7_, for growth. However, since the culture used in the experiments had a reduced microbiome rather than being fully axenic, it remains possible that the remaining bacteria supplied B_1_ or B_7_ and obscured *P. bahamense*’s requirement of those vitamins. These results are in line with previous genomic surveys [8] and bioassays [9], indicating that most HAB dinoflagellates are B_12_ auxotrophs. While *P. bahamense* was not included in these studies, Santos and Azana [46] found that maximum biomass tripled in antibiotic-treated cultures of *P. bahamense* var. *compressum* when the B-vitamin concentrations in f/2 media [47] were also tripled. Together, these results strengthen the case that B_12_ auxotrophy is a commonly shared trait among toxigenic dinoflagellates.

The *P. bahamense* strain used in this study required > 0.369 pM B_12_ to reach a chlorophyll fluorescence commensurate with 100,000 cells L^-1^, the minimum cell density designated by Florida state regulatory agencies as a bloom (Fig. 4, S4). The *P. bahamense* growth rate stabilized at 0.369 pM B_12_ and remained stable at higher B_12_ concentrations, even though higher B_12_ concentrations further increased the maximum chlorophyll fluorescence reached (Fig. 4). Similarly, the saxitoxin-producing dinoflagellates *Alexandrium pacificum* and *Gymnodinium catenatum* required > 0.34 and > 1. 52 pM B_12_, respectively, for measurable growth [8]. The brevetoxin-producing *Karenia brevis* requires > 0.01–1.0 pM B_12_ for measurable growth, depending on the strain tested [8, 9]. Thus, in addition to having an absolute B_12_ requirement, toxigenic dinoflagellates may have especially high B_12_ needs due to their unique physiologies.

Phytoplankton are generally considered auxotrophic for B_12_ if they lack the gene encoding B_12_-independent methionine synthase, but B_12_ may support dinoflagellate growth through other mechanisms as well [48]. Vitamin B_12_ is a cofactor for methylmalonyl-CoA mutase (MCM)—the enzyme that converts methylmalonyl-CoA to succinyl-CoA and is required for the breakdown of odd-chain fatty acids and several amino acids. The metabolites from MCM-catalyzed breakdown of amino acids are directed into the Krebs cycle, supporting energy production in eukaryotic cells [48, 49]. Moreover, S-adenosylmethionine (SAM), the activated form of methionine, acts as a universal methyl donor. SAM cycle methylation is key for regulating gene expression through DNA and histone methylation, and is also central to saxitoxin biosynthesis [8]. Given the large genomes characteristic of dinoflagellates [50], extensive methylation may be required for gene regulation [51], which would increase usage of the SAM cycle and elevate B_12_ demand.

### *Bacteria in seawater collected during a bloom supported* P. bahamense *growth*

Many studies have demonstrated that bacteria can fulfill phytoplankton B_12_ needs (e.g., [23, 36, 52]), including those of HAB-forming dinoflagellates (e.g., [17, 22]). Therefore, we tested whether bacteria recruited from several sources could rescue *P. bahamense* from B_12_ limitation. Only the bacterial community recruited from IRL seawater collected during a *P. bahamense* bloom was able to support *P. bahamense* growth in media without B_12_, although these cultures only reached about half the total biomass of cultures in replete media (Fig. 5). The maximum Chl-*a* fluorescence in these cultures was 1,140 RFU, just shy of 1,457 RFU—the value corresponding to the 100,000 cells L^-1^ bloom level (Fig. S4). Moreover, the growth rates in cultures with bacteria from IRL seawater were similar in replete media and media missing B_12_. Together, these results suggest that bacteria present during the *P. bahamense* bloom in IRL and transferred to our culture may supply B_12_ and support bloom development.

To evaluate which bacteria in the IRL seawater may provide B_12_, we identified taxa that were exclusively recruited from this source. Genera in the families Saprospiraceae (*Lewinella*) and Enterobacteriaceae (*Klebsiella* and *Enterobacter*) were consistently present in cultures with bacteria from IRL seawater and absent in cultures with bacteria from OTB seawater or IRL *P. bahamense* culture. *Lewinella* is not a known B_12_-producer, but is commonly associated with phytoplankton [53] and macroalgae [54], and may be particularly suited to degrading algal polysaccharides [53]. *Klebsiella* species are prolific B_12_ producers and are added to fermented foods for B_12_ enrichment [41]. *Enterobacter* species are similarly B_12_ producers [43]. In addition to being found in foods, *Klebsiella* and *Enterobacter* reside in mammalian small intestines [55] and are found in wastewater [44, 45]. *Alteromonas spp*., which are typically B_12_ auxotrophs [39], were most abundant in the smaller size fraction of communities recruited from IRL seawater when B_12_ was not included in the media, suggesting they may also benefit from B_12_ produced by Enterobacteraceae or other bacteria recruited from IRL. In contrast, Rhodobacteraceae genera—*Ruegeria* and *Sulfitobacter*—that are often found in phytoplankton cultures and associated with phytoplankton blooms [56] were found in all of our cultures and were not specific to those that grew without B_12_ added to media (Fig. 6, 7).

Surprisingly, bacteria recruited from the IRL culture did not support *P. bahamense* growth in the absence of B_12_, despite the IRL culture microbiome being fairly diverse and likely representing the seawater community when the culture was initiated. The diversity of culture microbiomes can diminish with culture age [56], and B_12_-producing bacteria may be outcompeted if the media is replete in B_12_. The OTB seawater may have originally hosted B_12_-producing bacteria that were lost in the extensive filtering used to remove eukaryotic phytoplankton from the inoculum, but it is also plausible that B_12_-producing bacteria were absent at the time of collection since *P. bahamense* was also absent. Notably, this study reveals that distinct communities were recruited to *P. bahamense* from three sources, suggesting that *P. bahamense* is opportunistic or a generalist in its associations with bacteria.

In some cases, bacteria associated with phytoplankton cultures or blooms compete for essential nutrients [36, 57, 58]. Here, B_12_-producing bacteria may have been present in the communities recruited from OTB seawater, but other community members outcompeted *P. bahamense* for the resource. For instance, *Caulobacter* can change morphology and motility to survive oligotrophic conditions, making them especially good competitors [59]. Caulobacterales were present in all recruited bacterial communities, but were most abundant in communities recruited from OTB seawater. Thus, bacterial competition for essential resources may have affected the growth of the *P. bahamense* grown with bacteria from OTB seawater.

### Conclusions & future directions

The results from this study demonstrate that *P. bahamense* is auxotrophic for B_12_ and that potentially anthropogenic bacteria present in the IRL may supply B_12_ to support bloom formation. Bacteria from OTB seawater did not support *P. bahamense* growth when B_12_ was withheld, but these bacteria were collected when *P. bahamense* was not detected in the environment. Future work should investigate B_12_ sources that support *P. bahamense* bloom formation by seeking to identify B_12_-producing bacteria that co-bloom with *P. bahamense* or determining if B_12_ is directly introduced to the ecosystem in wastewater. OTB and IRL are highly urbanized and receive excess nutrients from wastewater treatment plants [60], and the prolonged residence times [61] in these estuaries may also lead to an accumulation of B_12_. Given that increased B_12_ levels could exacerbate blooms, anthropogenic sources of B_12_ should be investigated in tandem with B_12_-synthesizing bacteria. Ultimately, measuring and tracing B_12_ in OTB and the IRL could disentangle the sources of B_12_ to these ecosystems and the relationship between anthropogenic B_12_ input and B_12_-producing bacteria with HABs. Such an effort may advance our ability to predict the severity of not just *P. bahamense* blooms, but other HABs as well.

## Acknowledgements

Research was funded by the University of South Florida College of Marine Science (USF CMS), with additional support from the Phycological Society of America Grants-in-Aid of Research and the Sanibel Captiva Shell Club grant. LRH was supported by the Abby Sallenger Endowed Fellowship in Marine Science and the Anne and Werner Von Rosenstiel Fellowship in Marine Science, awarded through USF CMS. We would like to thank the Florida Fish and Wildlife Conservation Commission (FWC) Fish and Wildlife Research Institute (FWRI) for *P. bahamense* culture loans and seawater sampling. Illumina metabarcode sequencing was performed by Georgia Genomics and Bioinformatics Core at the University of Georgia (GGBC, UG Athens, GA, RRID:SCR_010994).

## Author Contributions

MMB planned the project. LRH, LD, and MMB performed research. LRH and MMB analyzed data and wrote the manuscript. CL and SS reviewed and edited manuscript drafts.

## Data Availability

All sequence data from this project is available on NCBI SRA under PRJNA1315153. Code, data analysis pipelines, and raw data are available in a GitHub repository: https://github.com/lruggles1/Pyrodinium_bahamense_growth.git. The reproducible R Markdown analysis pipeline is further available as an interactive HTML file: https://lruggles1.github.io/Pyrodinium_bahamense_growth/pyro_growth_meta.html.

