## Supplemental Material for "B-vitamin requirements and phycosphere interactions for the HAB-forming dinoflagellate, *Pyrodinium bahamense* var. *bahamense*"

**Supplementary Material**

**Supplementary Tables**

**Table S1** Amplicon sequence variants (ASV) initially detected in the OTB *P. bahamense* culture

| Classification | ASV ID | Mean | Range | Also Found: (BLAST results) |
|---|---|---|---|---|
| O - *Rhizobiales*<br>F - *Hyphomicrobiaceae*<br>G - *Filomicrobium*<br>S - uncultured | 12a986 | 30% | 30–35% | Mangrove Sediment Samples<br><br>GenBank accession MH091208.1[1] |
| | e0169 | 9% | 7–13% | GenBank accession MH091208.1[1] |
| | aa0c6 | 0.07% | 0.04–1% | GenBank accession MH091208.1[1] |
| | f6a0f | 0.06% | 0.04–0.09 % | GenBank accession MH091208.1[1] |
| | ff08ec | 0.05% | 0–0.1% | GenBank accession EU488009.1[1] |
| O - *Rhizobiales*<br>F - *Hyphomicrobiaceae*<br>G - uncultured | c02844 | 1.1% | 0.1–2.6% | Date Palm rhizosphere, contaminated soils (*in situ* and experimental)<br><br>GenBank accession OK235762.1[2]<br><br>GenBank accession HQ697761.1[2]<br><br>GenBank accession KP098952.1[2] |

| | | | | |
|---|---|---|---|---|
| O - *Microtrichales*<br>F - uncultured | 07ae9a | 25.2% | 14–35% | Coastal wetland soil, alkaline soil, mine tailings, contaminated groundwater<br><br>GenBank accession KM840989.1[3]<br><br>GenBank accession HF558551.1[4]<br><br>GenBank accession JQ427269.1[5]<br><br>GenBank accession JQ665390.1[2] |
| | 9b8acc | 7.9% | 6.1–9.7% | GenBank accession KM840989.1[6] |
| O - *Oceanospirillales*<br>F - *Alcanivoraceae*<br>G - *Alcanivorax*<br>S - *Alcanivorax venustensis* | e9034 | 25% | 18–34% | Seamount in the Arabian Sea<br><br>GenBank accession PP516259.1[7] |

### Supplementary Figures

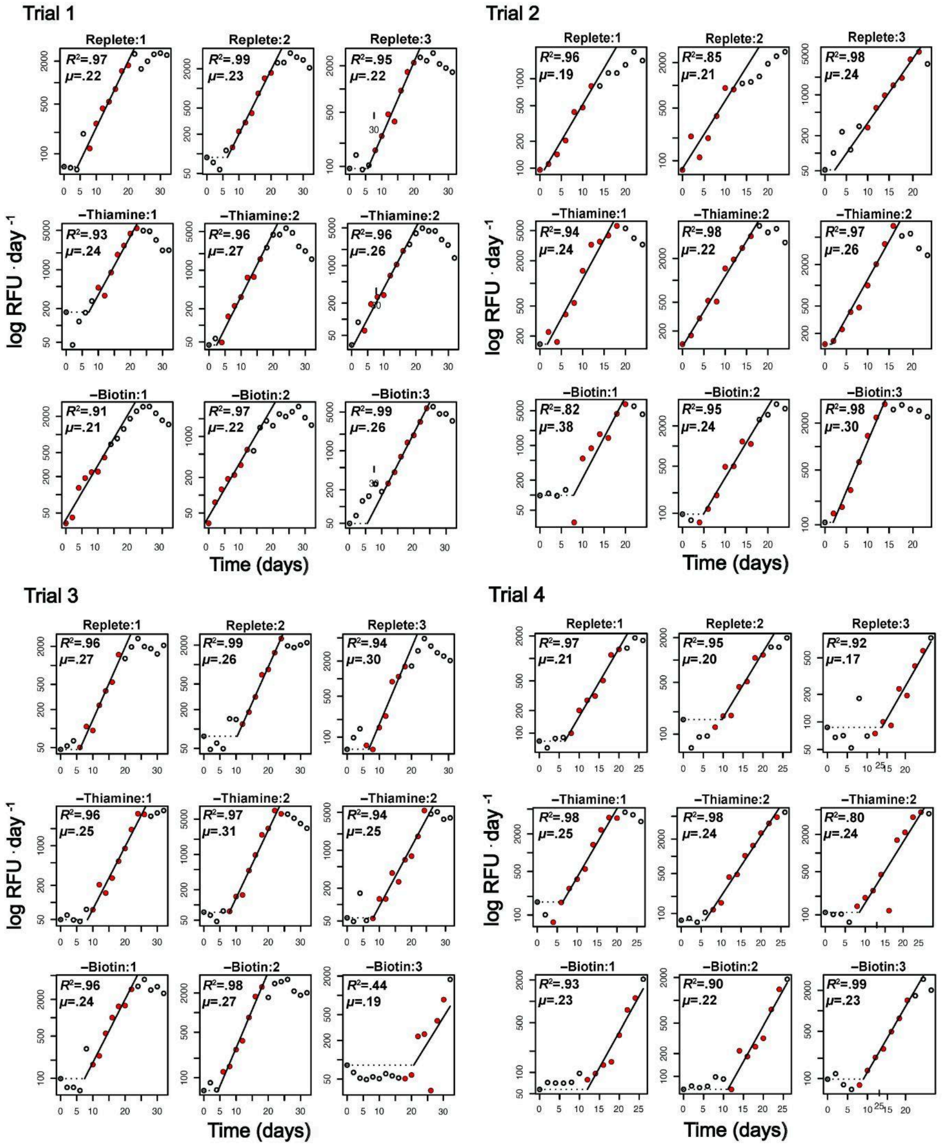


**Fig. S1** Growth curves for OTB *P. bahamense* (FWC1006) grown in replete L1/4, L1/4 –thiamine, and L1/4 –biotin media treatments for four trials. Red dots indicate the region of the curve used to calculate the specific growth rate ($\mu$; day$^1$), with black lines representing the line fitted using the all_easylinear function of the growthrates R package. Individual $\mu$ and $R^2$ are displayed on each plot for each replicate.

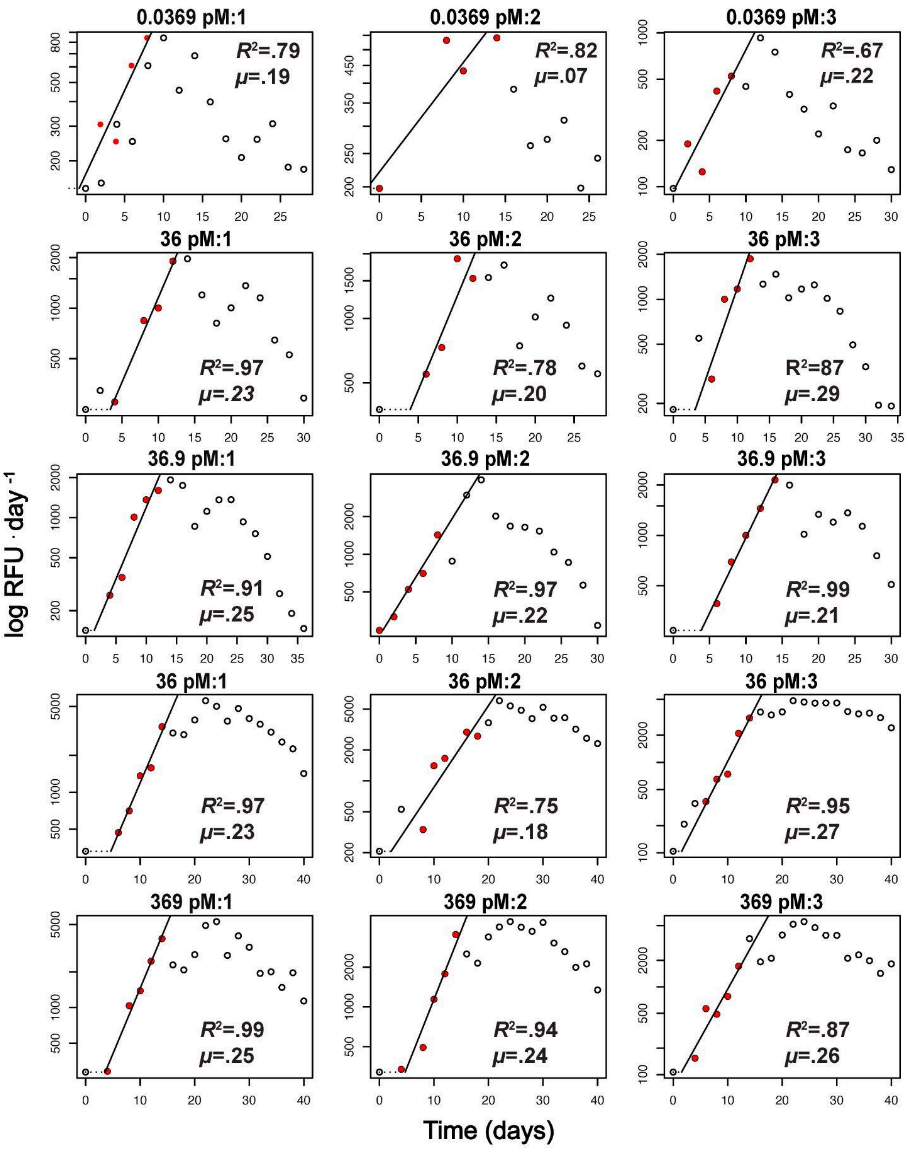


**Fig. S2** Growth curves for OTB *P. bahamense* (FWC1006) grown in five $B_{12}$ concentrations. Red dots indicate the region of the curve selected to calculate the specific growth rate ($\mu$; day¹), with black showing the line fitted using the all_easylinear function of the growthrates R package in RStudio. Individual $\mu$ and $R^2$ are displayed on each plot for each replicate. Note: slopes in 0.0369 pM of $B_{12}$ and replicate one in 0.369pM of $B_{12}$ were not significant.

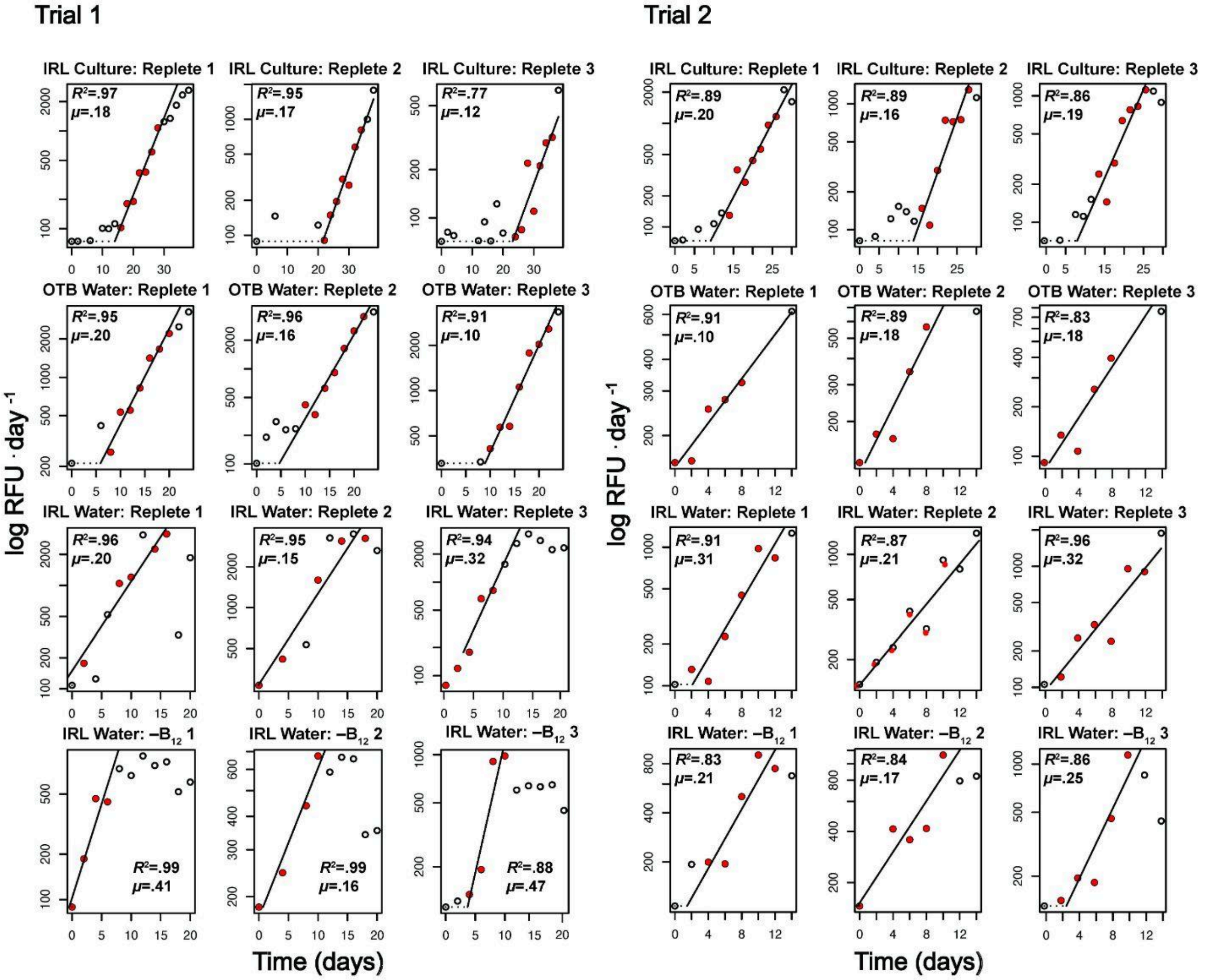


**Fig. S3** Growth curves of OTB *P. bahamense* (FWC1006) with bacteria from: IRL culture, OTB seawater, and IRL seawater. Cultures were grown in replete L1/4 media and L1/4 $-B_{12}$ media for two trials. Red dots indicate the region of the curve selected to calculate the specific growth rate ($\mu$; day$^{-1}$), with black lines indicating the line fitted using the all_easylinear function of the growthrates R package in RStudio. Individual $\mu$ and $R^2$ are displayed on each plot for each replicate.

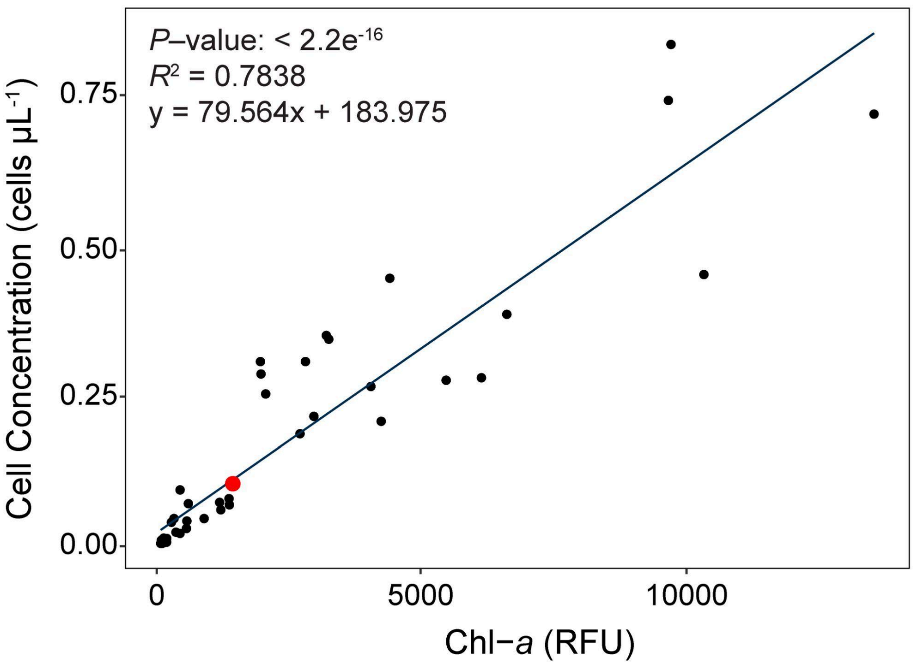


**Fig. S4** Linear regression of *P. bahamense* cell concentration (cell µL$^{-1}$) and *in vivo* Chlorophyll-*a* fluorescence (Chl-*a*) values. Triplicate biological replicates of *P. bahamense* were grown in L1/4 media made with sterile 25 PSU artificial seawater (ASW; Instant Ocean) at 24 °C on a 12:12 h light: dark cycle with 50% illumination (~115 µM photons m$^{-2}$ s$^{-1}$). *In vivo* Chlorophyll-*a* fluorescence (Chl-*a*) and imaging flow cytometry (ThermoFisher Attune CytPix) cell enumeration measurements were taken on triplicate technical replicates within each biological replicate every 48 h. The red point indicates bloom level (100,000 cells L$^{-1}$), which was calculated to be 1,457 RFU.

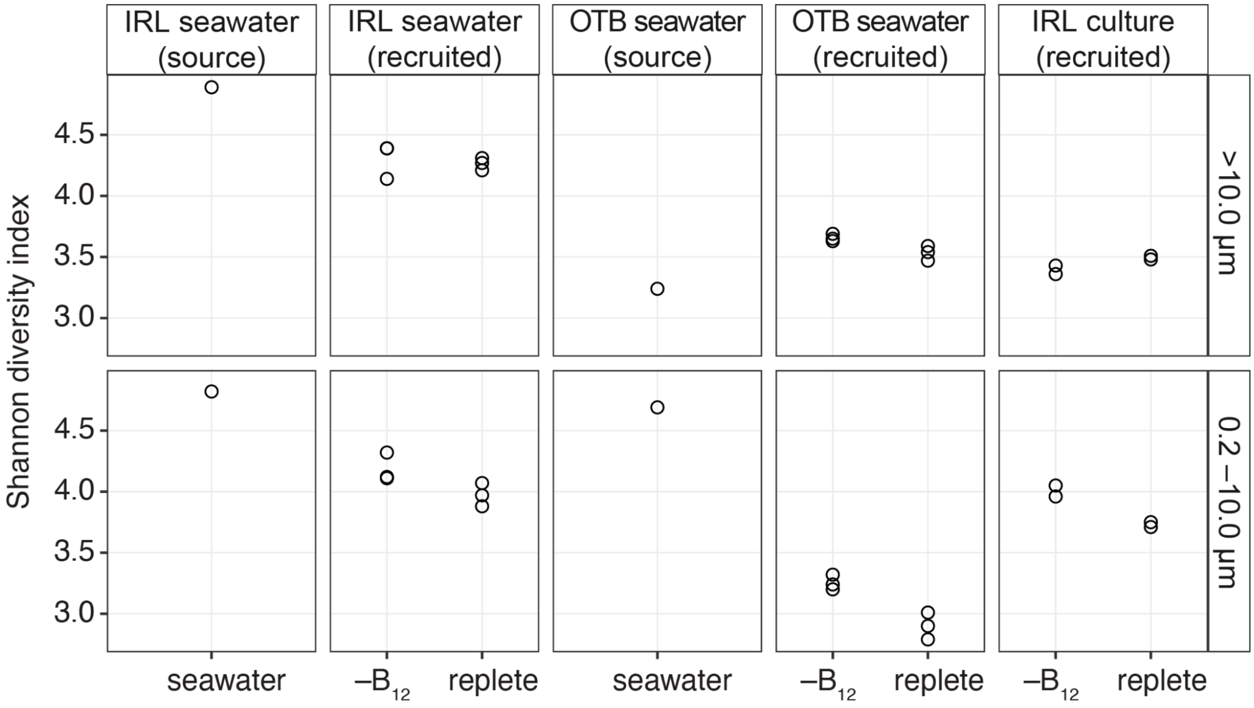


**Fig. S5** Shannon diversity indices for bacterial communities recruited by OTB *P. bahamense* from three sources (IRL Seawater, OTB seawater, IRL culture) and in the seawater sources. Shannon diversity was calculated in the EPI2ME 16S analysis pipeline.

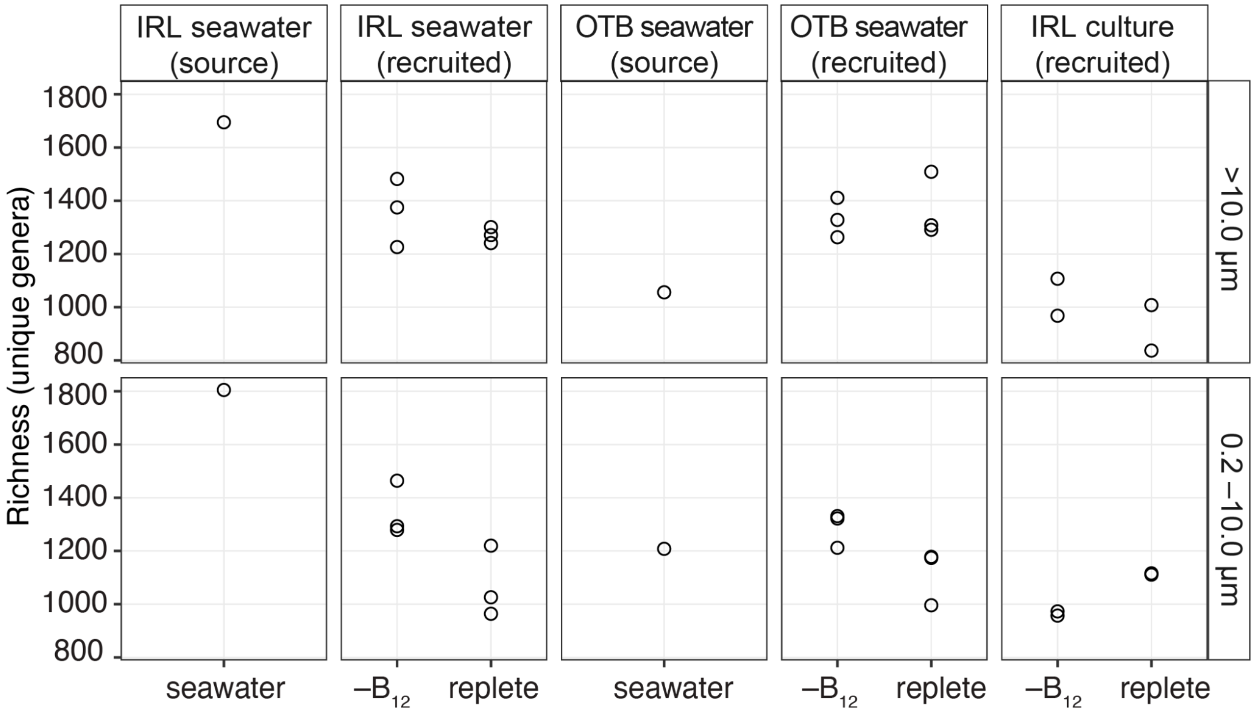


**Fig. S6** Richness (unique genera) for bacterial communities recruited by OTB *P. bahamense* from three sources (IRL Seawater, OTB seawater, IRL culture) and in the seawater sources. Richness was calculated in the EPI2ME 16S analysis pipeline.
